# To start or to discontinue the pill – changes in progestogens reflected by resting-state connectivity and positive mood

**DOI:** 10.1101/2022.09.21.508780

**Authors:** A-C.S. Kimmig, P. Friedrich, B. Drotleff, M. Lämmerhofer, I. Sundström-Poromaa, S. Weis, B. Derntl

**Affiliations:** Department of Psychiatry and Psychotherapy, Tübingen Center for Mental Health (TüCMH), University of Tübingen, Tübingen, Germany; Graduate Training Centre of Neuroscience, International Max Planck Research School, University of Tübingen, Tübingen, Germany; Institute of Neuroscience and Medicine, Brain & Behaviour (INM-7), Research Centre Jülich, Jülich, Germany; Institute of Systems Neuroscience, Medical Faculty, Heinrich Heine University Düsseldorf, Düsseldorf, Germany; Institute of Pharmaceutical Sciences, University of Tübingen, Tübingen, Germany; Department of Women’s and Children’s Health, Uppsala University, Uppsala, Sweden; LEAD Graduate School and Research Network, University of Tübingen, Tübingen, Germany

## Abstract

Oral contraceptive (OC) intake has been associated with alterations in functional brain architecture and socio-affective processes. However, most previous studies have been limited by cross-sectional designs and/or did not account for synthetic sex hormone concentrations. The aim of this longitudinal study was to determine the effects of starting vs discontinuing OCs on socio-affective functions such as mood and emotion cognition, and to identify their possible neuroendocrinological substrates.

To this end, 88 young healthy women performed the behavioral and fMRI measures twice, three to eight months apart: 26 natural cycling women twice during menstruation, 26 OC users twice during OC intake, 25 OC discontinuers and 11 OC starters before and after discontinuation or start, respectively. In addition to mean-based analyses, we used intersubject representational similarity analyses to determine relationships between interindividual variability in within-subject changes of hormone profiles, including concentrations of endogenous and synthetic hormones, region-specific resting state functional connectivity (parcelwise RSFC) and socio-affective measures.

Across the whole sample, interindividual patterns of changes in RSFC of fronto-parietal regions, parts of the left hippocampus and the right cerebellum reflected change patterns of progestogen levels. For the right superior orbitofrontal gyrus (OFG), a trinity of idiosyncratic patterns was found in changes of progestogens, RSFC and positive mood. Active OC intake was associated with higher self-reported depressive symptoms in OC discontinuers (and starters). Emotion recognition performance was not associated with changes in hormone profiles or RSFC.

Overall, progestogens rather than estrogens appear to be associated with functional brain architecture of the frontal and subcortical/cerebellar regions and positive mood. The right superior OFG represents a possible neural substrate for progestogen-induced changes in positive mood. This study indicates the importance of a multidimensional, longitudinal approach when being interested in effects of hormonal contraception on women’s brain and behavior.

---

Estradiol and progesterone are the predominant ovarian hormones involved in the menstrual cycle, modulating female fertility during reproductive years. Apart from their local binding sites in the ovaries (e.g., regulating follicular growth, endometrial buildup, and shedding), they also bind to several receptors in the brain after crossing the blood-brain barrier[1]. Besides the hypothalamus, dense sex hormone receptor sites in the brain are located within limbic areas including the amygdala and hippocampus, as well as frontal and cerebellar regions[2-4]. Neural functions, including functional connectivity, and morphology are influenced by sex hormones through mechanisms such as synaptic pruning, neurite outgrowth, dendritic branching, and myelination[5-8]. Worldwide more than 150 million women use oral contraceptives (OCs) for birth control[9]. Through negative feedback on the hypothalamus and pituitary gland, OCs suppress hormonal fluctuations, disrupting the menstrual cycle. Similar to endogenous sex hormones, the synthetic hormones (i.e., ethinylestradiol and progestin) contained in the OCs exert estrogenic, progestogenic, and androgenic effects on hormone receptors spread over the body, including the brain[10, 11]. Therefore, not only interaction with natural hormonal fluctuations, but also additional binding of the synthetic compounds to hormone receptors in the brain could lead to functional and morphological neural changes, especially in brain regions involved in socio-affective processing[12]. Indeed, both menstrual cycle phase and OC intake have been associated with altered socio-affective processes, including mood, fear processing, emotion recognition, and empathy as well as sexual behavior[12-16]. Given the high prevalence of OC use and its potential psychological side effects, it is critical to investigate how functional brain architecture is influenced by neuroendocrinological mechanisms.

Resting-state (i.e., task-unrelated) functional connectivity (RSFC) is thought to capture the coupling dynamics within and between large-scale brain networks[17-19] and thus provides insight into intrinsic brain architecture. It is increasingly used to investigate, amongst others, the neurobiological basis of behavior as well as socio-affective processes[20]. Functional connectivity refers to the extent to which the neural activity of two brain regions covaries over time. In terms of neuroendocrinological mechanisms, there is preliminary evidence that cortico-cortical, as well as cortico-subcortical RSFC, is affected by fluctuating sex hormones during the menstrual cycle and by OC intake. A comprehensive review article reported that the menstrual cycle most consistently modulates connectivity of the middle frontal gyrus (MFG) and the inferior parietal lobule[21]. RSFC between the MFG and regions of the executive control network was reduced in OC users compared to follicular naturally cycling (NC) women[22]. Moreover, studies with dense-sampling of single-subjects or longitudinal design, suggest hormone-related reorganization of functional brain networks across the menstrual cycle[23-26] as well as OC intake phase[27] and with the start of OC intake[28, 29], with the latter consistently leading to relatively reduced amygdala and (para-)hippocampal RSFC.

The hormone-induced modulation of functional brain architecture involved in socio-affective processing, e.g. the MFG, the amygdala, and the (para-)hippocampus[30], could be potential neural correlates of the previously identified hormone-related differences in a variety of self- and other-related emotional processes[12]. To date, mood and emotion recognition are among the most studied socio-affective processes related to the menstrual cycle and OC intake, probably due to their high relevance to social functioning and mental health[31-33]. Although no direct link has been investigated, women who had reduced amygdala and (para-)hippocampal RSFC after starting OC intake also reported a flattening of positive mood[28]. Moreover, severe OC-induced depressive symptoms are thought to affect only a subset of hormone-sensitive women[31, 32], presumably due to an altered sensitivity of the GABAergic system via progestin induced changes in GABA-A receptor subunit expression[34], which likely affects functional brain architecture[35]. Hence, altered hormone-related RSFC may indeed be part of the neural mechanisms underlying hormone-related mood changes. Similarly, OC intake appears to modulate the relationship between amygdala RSFC and emotion recognition performance differently than the menstrual cycle[36]. Therefore, hormonal influences on brain regions involved in emotion recognition may lead to hormone-related differences in emotion recognition performance. In line, a recent review suggested that OC intake appears to be associated with reduced emotion recognition performance[33]. In addition, differences in performance have been observed in NC women in the follicular phase compared to the luteal phase. While endogenous estradiol generally exerts positive effects and progesterone shows a bimodal (i.e., inverted U-shape) association with mood[37, 38], no consistent relationships have been found between endogenous sex hormones and emotion recognition. The interaction between endogenous and synthetic sex hormones on the functional architecture of the human brain, and consequently on socio-affective processes such as mood and emotion recognition, is not known, as studies do not usually examine circulating synthetic sex hormones.

Overall, endogenous and synthetic sex hormones seem to modulate the functional organization of the brain, especially in limbic and frontal brain regions, which have a high density of hormone receptors and are also involved in socio-affective processes such as mood and emotion recognition. However, most of the findings to date are based on cross-sectional studies, making causal inferences difficult. Initial longitudinal studies examining the effects of OC onset on the brain reported changes in brain anatomy and functional organization[27-29]. Yet not a single study so far investigated the reversibility of these OC-induced effects on the brain and related socio-affective processes by directly examining OC discontinuation. Furthermore, the fact that synthetic sex hormone concentrations have not been considered in previous studies further complicates the understanding of OC-induced effects on the brain and socio-affective processes. Given the high prevalence of OC intake and its potential impact on female mental health, it is crucial to systematically investigate the potential effects of OCs on women’s socio-affective functions such as mood and emotion recognition, and the neuroendocrinological mechanisms that may underlie them. To this end, we conducted a longitudinal study in which women’s endogenous and synthetic sex hormones, RSFC, mood, and emotion recognition performance were measured twice over three to eight months (see Figure 1).

**Figure 1.**
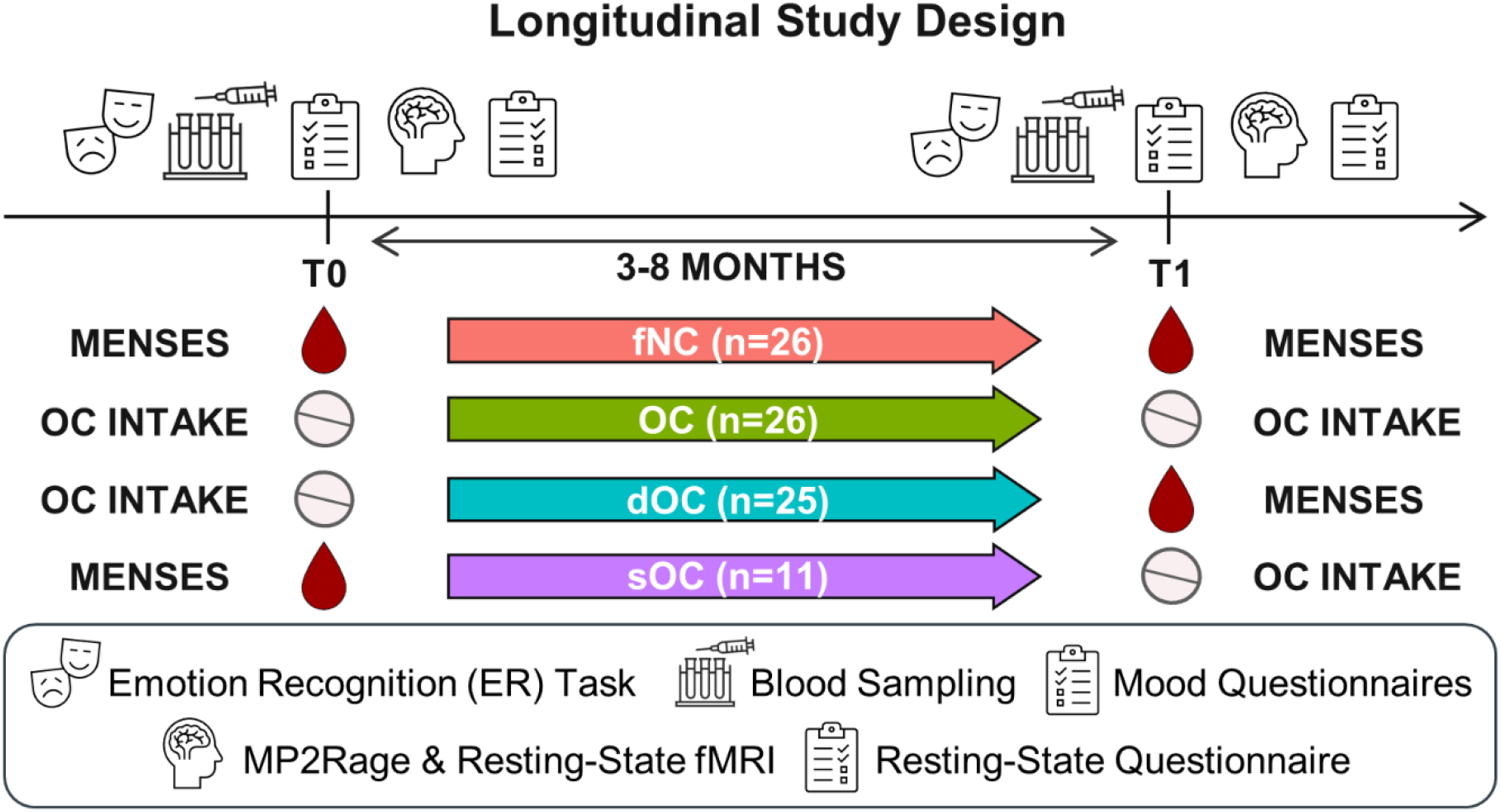
Overview of Longitudinal Study Design. Participants were measured twice over three to eight months, either both times during 1) menses (i.e., fNC group), 2) active OC intake (i.e., OC group), or before and after 3) discontinuation of OC intake (i.e., dOC group), or 4) start of OC intake (i.e., sOC group). Hormonal, neural (functional and structural MRI) and behavioral data (measures of mood and emotion recognition) were collected at both time points. fNC – early follicular naturally cycling, OC – continued oral contraceptive intake, dOC – oral contraceptive discontinuation, sOC – oral contraceptive start.

We included young, healthy women with relatively stable hormonal profiles (i.e., both time points measured during the early follicular or active OC intake phase) and women with changing hormonal profiles (i.e., women starting or discontinuing OC intake) to determine how

(1) stability or change of OC status/hormonal profile affects a) hormone concentrations, b) mood measures including positive and negative mood as well as depressive symptoms, and c) overall facial emotion recognition performance analyzed by group comparisons,

and more specifically whether

(2) patterns of change in sex hormone (i.e., estrogens and progestogens) concentrations resemble changes in a) locally specific RSFC, b) mood-related measures, and c) overall facial emotion recognition performance, and
(3) patterns of change in a) mood-related measures and b) emotion recognition performance are reflected in changes in hormone-modulated locally specific RSFC.

Locally specific RSFC was determined using a whole-brain parcellation approach in which the brain was divided into 268 functionally similar parcels[39] and RSFC was averaged across all voxels in a parcel. The two latter research questions were investigated using intersubject representational similarity analyses (IS-RSA). IS-RSA is a method of analysis that allows individual differences across modalities (i.e. hormonal, RSFC, and socio-affective measures) to be linked and to determine whether individuals who are more similar in one modality (e.g. change in hormones) are also more similar in another modality (e.g. change in parcelwise RSFC) using second-order isomorphisms[40, 41]. To this end, each subject’s unique pattern of synchrony with other subjects is used to form a covariance matrix (i.e., a representative distance matrix, RDM) per modality. Subsequently, the patterns of inter-subject synchrony between modalities are compared by correlating them with each other. The advantage of using second order isomorphisms is that the geometry of two modalities, rather than their physically different quantities, can be compared. Furthermore, IS-RSA complements mean-based group comparisons as it is more sensitive to interindividual variance and allows for continuous analysis across the entire sample[40]. Lastly, estradiol and progesterone have been suggested to exert individual, hormone-specific effects on RSFC that depend on brain localization as well as hormone status and hormone fluctuations[21, 42]. To better approximate the individual roles of estrogens (i.e., estradiol and ethinylestradiol) and progestogens (i.e., progesterone and progestin) in RSFC and socio-affective functions, potential synchronizations between changes in hormone profiles, parcelwise RSFC and mood-as well as emotion recognition-related measures were therefore assessed separately for the different sex hormones using IS-RSA (see Figure 2).

**Figure 2.**
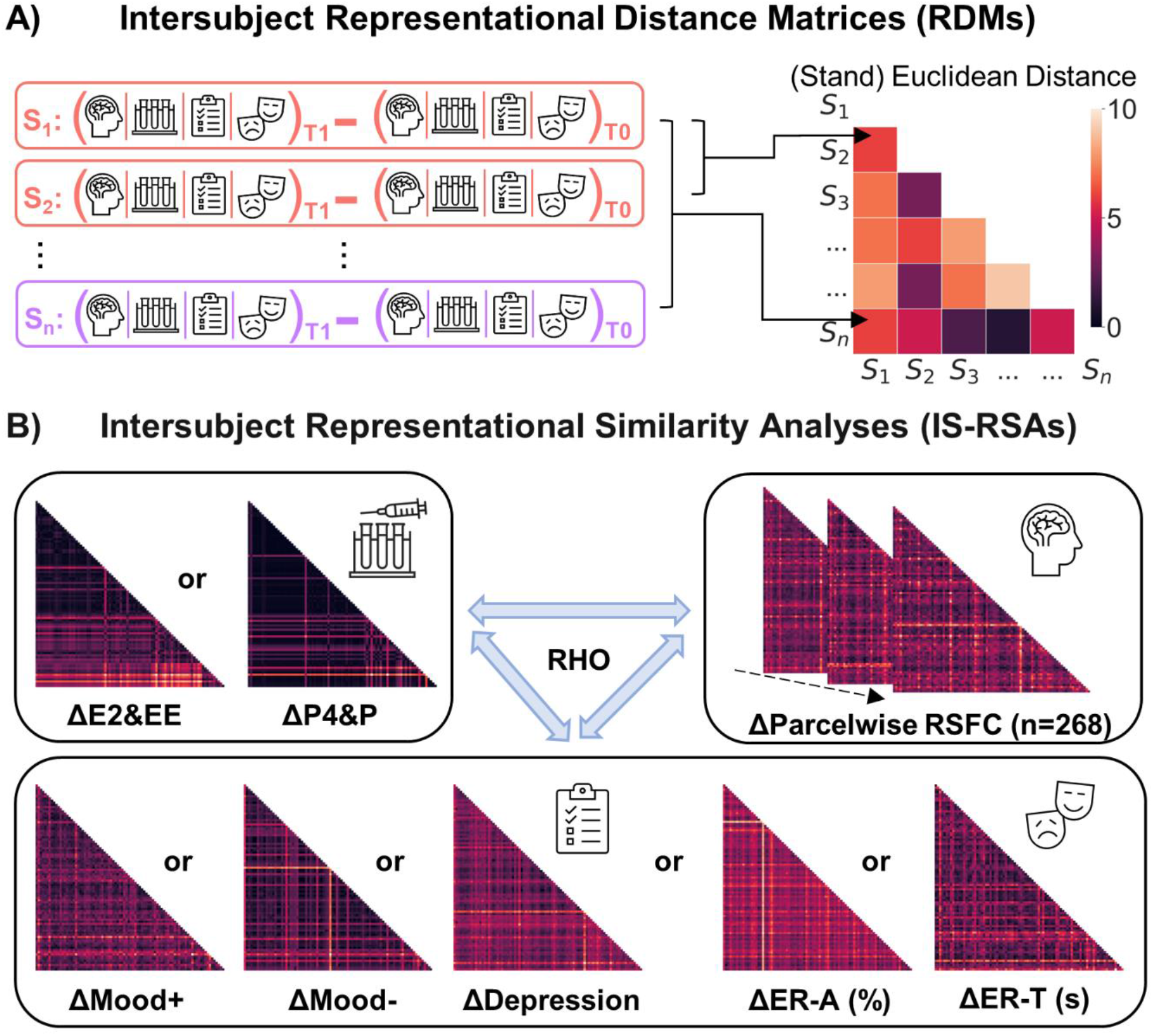
Representational Similarity Analysis Design. A) In addition to group differences based on means or medians, we were also interested in whether the patterns of change in the different measures were associated with each other. For this purpose, representational intersubject dissimilarity matrices (IS-RDMs) were calculated from the differences between T1 and T0 of each measure individually. B) The patterns of the IS-RDMs were then compared with permutation-based intersubject representational similarity analyses (IS-RSAs) using Spearman Rho. For the IS-RSAs with parcelwise RSFC (n=268 parcels), permutation-based multiple comparison correction was performed. Δ – T1 minus T0, E2 – Estradiol, EE – Ethinylestradiol, P4 – Progesterone, P – Progestin, RSFC – Resting-State Functional Connectivity, Mood+ – Positive Mood, Mood-– Negative Mood, ER-A – Emotion Recognition Accuracy, ER-T – Emotion Recognition Response Times

## Results

### Oral contraceptive intake impacts endogenous and synthetic sex hormone levels

The hormone concentrations of women with stable hormone profiles (i.e., women in the early follicular phase [fNC] or with continued OC use [OC]) and with changing OC status (i.e., starting [sOC] or discontinuing OC intake [dOC]) were analyzed with mixed ANOVAs and parametric or non-parametric post-hoc tests depending on data distribution. As expected, the presence of synthetic hormones significantly suppressed endogenous estradiol and progesterone concentrations, which reversed when OC intake was discontinued (group-by-time interaction: all F≥3.67, all *p*≤.015, all *η*^*2*^≥.12; see Table 1 and Section S1 in the Supplementary Materials for more detailed information on the post-hoc tests). The changes in estradiol (E2), ethinylestradiol (EE), progesterone (P4), progestins (P), and testosterone (T) between measurements are shown in Figure 3A. Synthetic hormones were no longer detected three to eight months after OC discontinuation. While synthetic P levels did not differ between the different groups at active OC intake phases (all *p*≥.346), synthetic EE levels were significantly higher in OC starters compared to OC users with continued intake (*H*(1)=13.48, *p<*.*001*). Contrary to our expectation of a stable hormone profile, OC users with continued OC intake showed a significant decrease in EE levels (*Z*=-2.31, *p*=.021). P levels, on the other hand, remained similar (all *p*≥.366). T levels were not altered by OC intake (all *p*≥.138), but the fNC group had a significant reduction in T concentrations at the second measurement (*t*(25)=3.16, *p*=.024).

**Figure 3.**
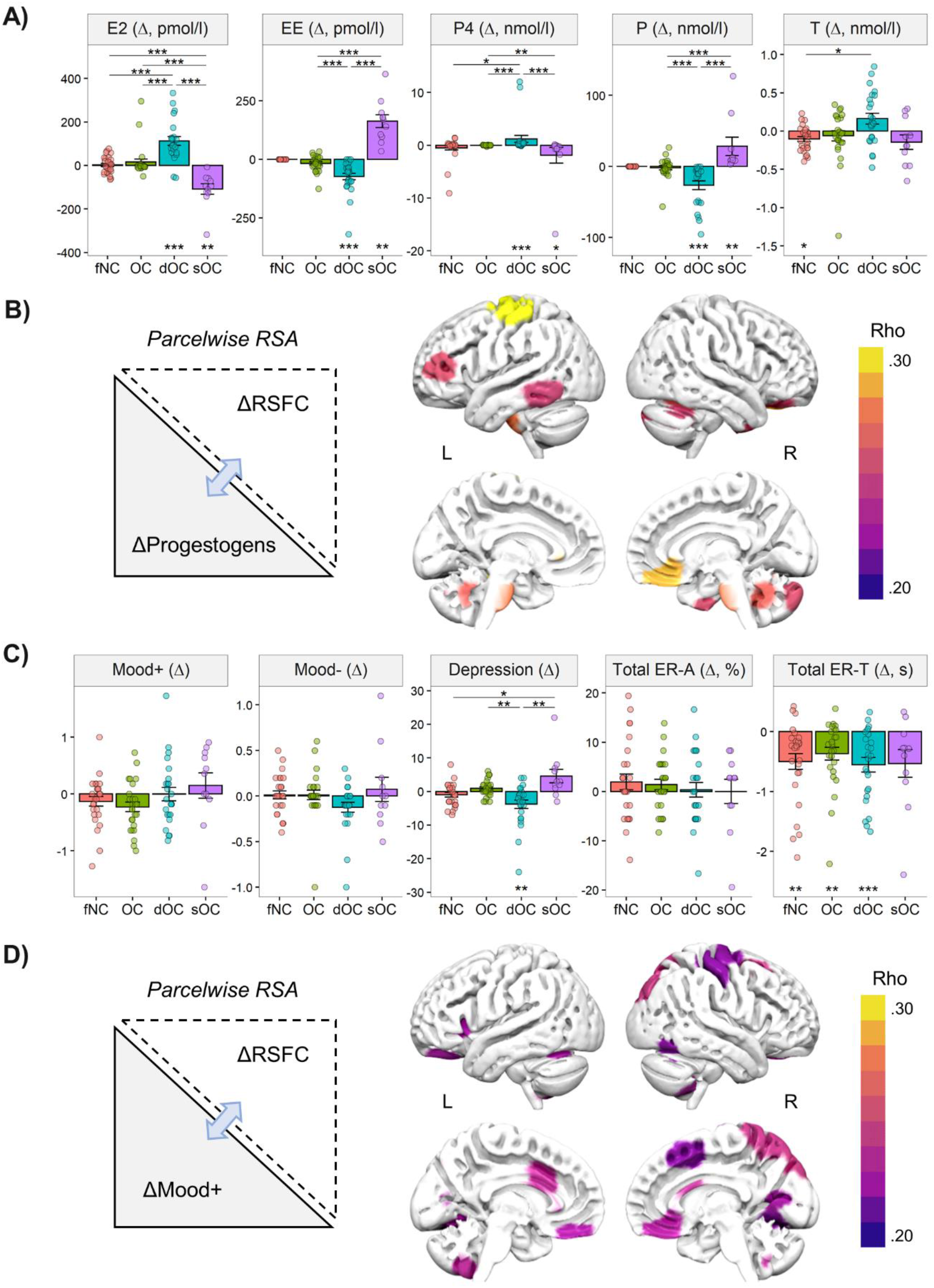
Group differences in hormonal and behavioral profiles and significant brain-hormone and brain-behavior associations. A) Bar charts illustrating the hormonal changes (ΔT1-T0) of estradiol (E), ethinylestradiol (EE), progesterone (P4), progestins (P), and testosterone (T) concentrations per group (i.e., fNC – red, OC – green, dOC turquoise and sOC – purple). B) Permutation-based multiple comparisons corrected significant parcels of IS-RSA between change patterns of parcelwise RSFC and progestogen concentrations (ΔP4 and P). Representative distance matrices (RDMs) were computed with Euclidean and standardized Euclidean distances, respectively. Significant Spearman’s Rho used to compare similarity between RDMs ranged between .25 and .30. Due to limitations of the surface-based brain graphic, the left hippocampus could not be visualized (see Table 2 for a complete list of significant parcels) C) Bar charts illustrating the behavioral changes (ΔT1-T0) of positive mood (Mood+), negative mood (Mood-), depressive symptoms (Depression), emotion recognition accuracy (ER-A), and emotion recognition time (ER-T) per group (i.e., fNC – red, OC – green, dOC – turquoise and sOC – purple). D) Permutation-based multiple comparisons corrected significant parcels of IS-RSA between change patterns of parcelwise RSFC and positive mood items (ΔMood+). Significance (*) annotations above bar charts represent differences in the change of concentrations from T0 to T1 between groups, whereas annotations below the bars refer to significant concentration changes within the respective group. Bonferroni correction was used to correct for multiple comparisons. Δ – T1 minus T0, E2 – Estradiol, EE – Ethinylestradiol, P4 – Progesterone, P – Progestin, RSFC – Resting-State Functional Connectivity, Mood+ – Positive Mood, Mood-– Negative Mood, ER-A – Emotion Recognition Accuracy, ER-T – Emotion Recognition Response Times, RSA – Representational Similarity Analysis, L - left, R – right. *p<.05, **p<.01, ***p<.001

**Table 1.**
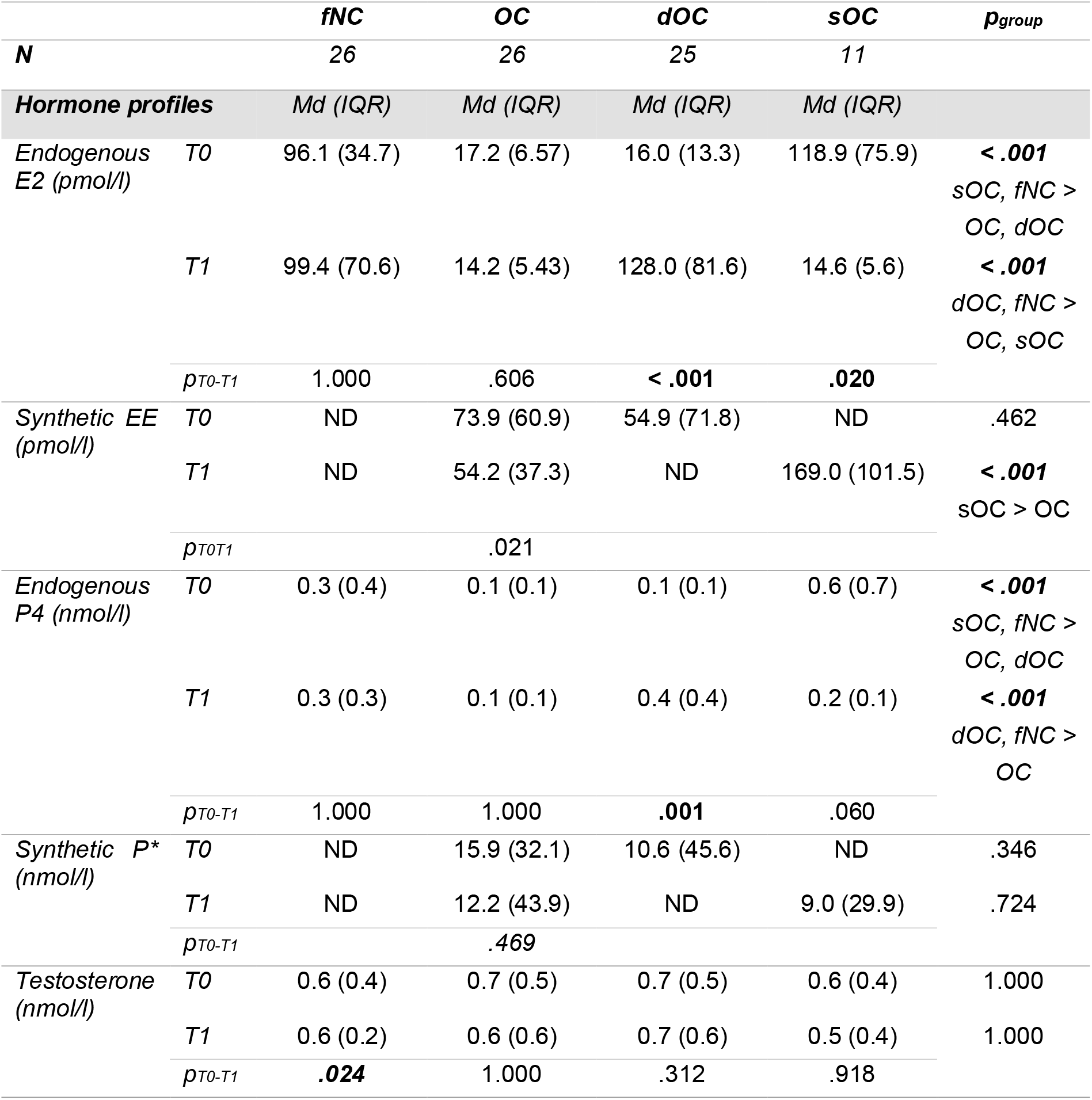
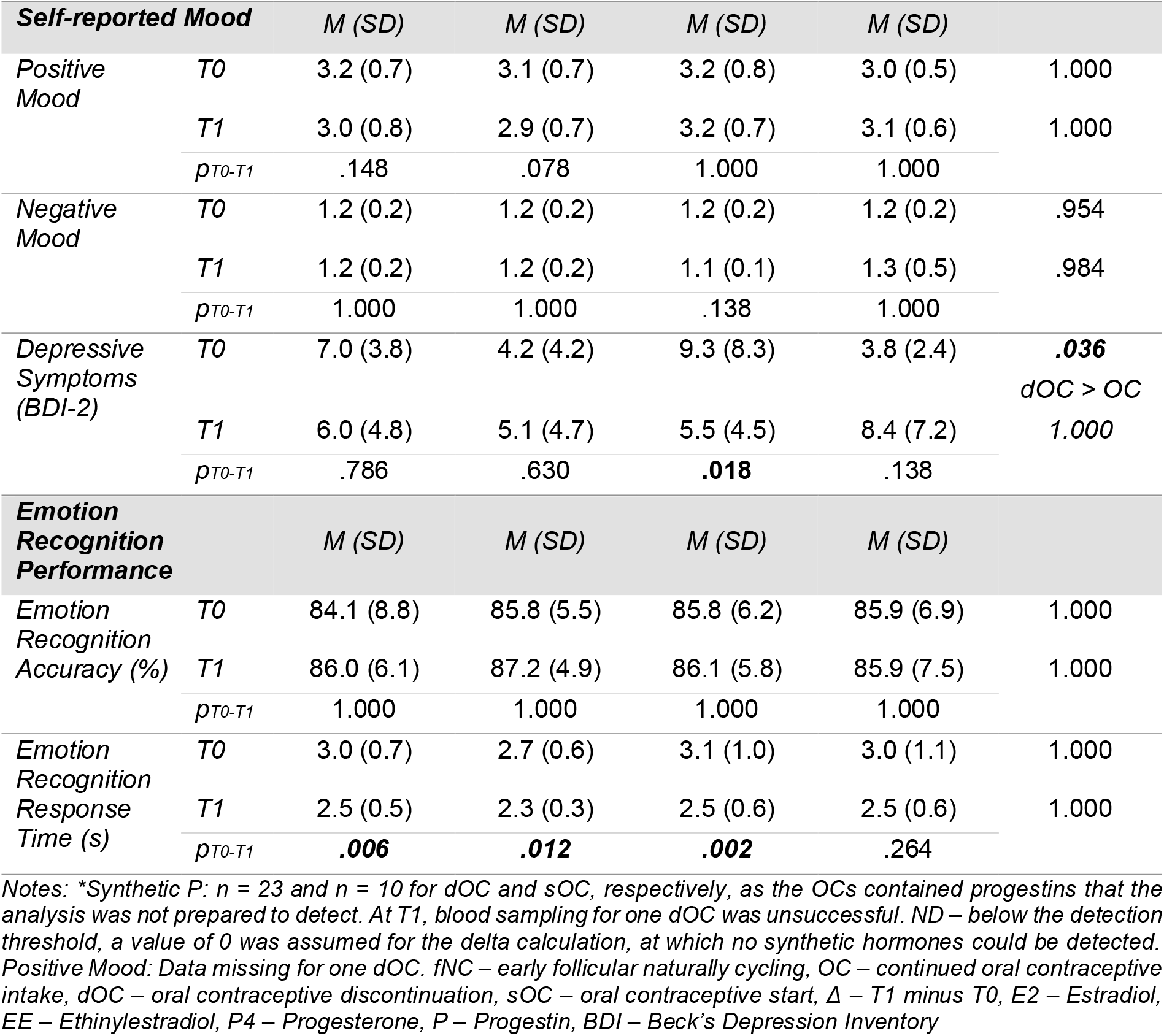
Multiple comparisons corrected contrasts of hormone levels (median (Md) and interquartile range (IQR)), self-reported mood and emotion recognition performance (percent correct as well as response times; mean (M) and standard deviation (SD)) per group and across the two measurement time points (T0, T1)

### Change patterns in hormonal profiles of progestogens, but not estrogens, are associated with changes in parcelwise RSFC

In the next step, we aimed to identify neural modulations of changes in estrogens and progestogens. To this end, we computed two separate inter-subject representative similarity analyses (IS-RSAs) to test whether change patterns across time points in estrogen and/or progestogen concentrations were reflected in the change patterns of parcelwise RSFC (see Figure 2).

**Table 2.** Overview of significant IS-RSA results (corrected for multiple comparisons) between change patterns of parcelwise RSFC and sex hormone profiles, mood parameters, and emotion recognition performance. Parcelwise MNI coordinates (center of mass) and functional network belonging are reported according to Finn, Shen [43].

| Shen Parcels | rho | MNI coordinates |  |  | Network | p<sub>corr</sub> |
| --- | --- | --- | --- | --- | --- | --- |
|  |  | x | y | z |  |  |
| Associated with progestogens (n=84 participants) |  |  |  |  |  |  |
| L PoCG | 0.30 | -35.9 | -23.3 | 65.6 | Motor | .009 |
| L Hippocampus | 0.29 | -35.7 | -24.8 | -14.9 | Subcortical-Cerebellar | .012 |
| R OFG | 0.29 | 5.1 | 34.9 | -17.4 | Default Mode | .016 |
| Brainstem | 0.28 | -6.1 | -18.9 | -36.8 | Subcortical-Cerebellar | .023 |
| R Cerebellum IX | 0.27 | 7.1 | -53.7 | -34.4 | Subcortical-Cerebellar | .032 |
| R Cerebellum VI | 0.26 | 21.1 | -54.8 | -23.8 | Subcortical-Cerebellar | .035 |
| L MFG/IFG | 0.26 | -43.0 | 42.0 | 11.0 | Frontoparietal | .041 |
| R Cerebellum Crus 2 | 0.26 | 11.7 | -84.1 | -34.6 | Frontoparietal | .041 |
| R FFG/ITG | 0.26 | 31.5 | 0.7 | -44.4 | Motor | .042 |
| R Superior OFG | 0.26 | 15.6 | 34.1 | -22.6 | Frontoparietal | .044 |
| R Cerebellum Crus 1 | 0.25 | 36.7 | -57.1 | -32.8 | Subcortical-Cerebellar | .047 |
| L FFG/ITG | 0.25 | -60.3 | -50.0 | -14.0 | Frontoparietal | .048 |
| Associated with positive mood (n=85 participants) |  |  |  |  |  |  |
| R Superior OFG | 0.25 | 18.9 | -81.8 | 41.5 | Visual I | .019 |
| R SFG | 0.25 | 25.2 | 12.4 | 49.4 | Frontoparietal | .021 |
| R Precuneus | 0.24 | 7.5 | -57.3 | 61.8 | Subcortical-Cerebellar | .022 |
| *L MCC* | 0.24 | -5.1 | 13.2 | 28.7 | Subcortical-Cerebellar | .028 |
| *L Cerebellum IX* | 0.24 | -8.7 | -55.2 | -52.1 | Default Mode | .030 |
| *R OFG* | 0.23 | 5.1 | 34.9 | -17.4 | Default Mode | .036 |
| *R Cerebellum VIII* | 0.23 | 33.8 | -47.2 | -53.9 | Subcortical-Cerebellar | .039 |
| *L FFG* | 0.23 | -25.9 | -63.1 | -12.3 | Visual I | .039 |
| *L OFG* | 0.23 | -8.2 | 39.7 | -21.4 | Medial Frontal | .040 |
| *L Insula Lobe* | 0.23 | -32.5 | 22.1 | 5.8 | Subcortical-Cerebellar | .041 |
| *R Calcarine Cortex* | 0.22 | 14.6 | -68.3 | 8.3 | Visual I | .041 |
| *R PreCG* | 0.22 | 26.3 | -12.9 | 66.2 | Motor | .049 |
| *R Lingual Gyrus* | 0.22 | 21.0 | -63.7 | -9.0 | Visual I | .049 |
| *R FFG/ITG* | 0.22 | 46.5 | -59.9 | -14.8 | Visual Association | .049 |
| *R PMFG* | 0.22 | 6.1 | 14.0 | 48.7 | Subcortical-Cerebellar | .050 |
| *R PoCG* | 0.22 | 42.0 | -23.4 | 53.4 | Motor | .050 |

Notes: All *p* values are corrected for multiple comparisons by permutation-based testing. L – left, R – right, PoCG – postcentral gyrus, OFG – orbitofrontal gyrus, MFG/IFG – middle frontal gyrus/inferior frontal gyrus, FFG/ITG – fusiform gyrus/inferior temporal gyrus, SFG – superior frontal gyrus, MCC – midcingulate cortex, PreCG – precentral gyrus, PMFG – posterior-medial frontal gyrus

Interindividual variation in the changes of progestogen hormonal profiles (i.e., RDM: standardized Euclidean distances for ΔP4 and ΔP) was positively reflected in brain parcels mainly belonging to the subcortical-cerebellum and frontoparietal networks[43] after permutation-based multiple comparison correction (see Figure 3B and Table 2 for IS-RSA results). Among parcels in the subcortical-cerebellum network, the highest synchrony with hormonal change was found in the left hippocampus, the brainstem, and three regions in the posterior lobe of the right cerebellum namely in lobule IX, lobule VI, and Crus1. With regards to parcels of the frontoparietal network, variation in the progestogen level changes were significantly resembled by the RSFC in the left MFG/IFG, the right cerebellum Crus2, the right superior OFG, and the left fusiform/inferior temporal gyrus (FFG/ITG). Furthermore, participants who were less similar in the changes of progestogen levels were accordingly also more dissimilar in their changes in RSFC of the left postcentral gyrus (PoCG), the right FFG/ITG, and the right OFG (see Table 2 for all test statistics and MNI coordinates).

After permutation-based correction for multiple comparisons, the IS-RSA of the change in estrogens (i.e., standardized Euclidean distances for ΔE2 and ΔEE) across the entire sample revealed no synchrony with the parcelwise RSFC change patterns (i.e., Euclidean distances for ΔRSFC per parcel; all *rho(86)*≤0.14, all *p*≥.235).

### OC discontinuers and starters report heightened self-reported depressive symptoms during OC intake, other measures of mood and emotion recognition are unaffected

To determine possible mean-based effects of initiation or discontinuation of OC intake on socio-affective functions such as positive and negative mood, depressive symptoms, and emotion recognition performance, 4 (between: group) by 2 (within: time points) mixed ANOVAs were performed for each behavioral measure. Table 1 provides an overview of all behavioral means separately for timepoint and for each hormone group. The mean change scores of the behavioral measures are depicted in Figure 3C.

While positive and negative mood showed no differences between the hormonal profile groups (group: *F*_*Mood+*_(3,83)=0.60, *p*=.616; *H*_*Mood-*_(3)=4.58, *p*=.206) and were not affected by time of measurement (group-by-time: all *F*≤1.70, all *p*≥.173), active OC intake had a significant effect on depressive symptoms in OC starters and discontinuers (group-by-time: *F*(3,84)=9.55, *p*<.001, *η*^*2*^=.25). OC discontinuers experienced a significant reduction (*Z*=-2.96, *p*=.018) of comparatively high depressive symptoms scores at T0 (*H*(3)=12.60, *p*=.036, dOC > OC users: *p*=.026), whereas no significant changes were observed within the other groups (all *p*≥.138). Using change scores (T1-T0, see Figure 3C), non-parametric post-hoc tests suggest that the change in depressive symptoms was significantly different in OC discontinuers compared to OC starters *(p*<.001) and OC users (*p*=.003). Furthermore, the increase in depressive symptoms in OC starters was significantly larger than in the fNC group (*p*=.014). There were no main effects of group (*H*(3)=5.02, *p*=.171) nor time (*F*(3,84)=0.09, *p*=.765).

For emotion recognition accuracy (i.e., the overall percentage of correctly identified trials, ER-A) or emotion recognition response times (i.e., mean response times of correctly identified trials, ER-T), no effects of start or discontinuation of OCs in contrast to women with a stable hormonal profile were found (group-by-time interaction: *F*_*ER-A*_(3,84)=0.31, *p*=.820 and *F*_ER-T_(3,84)=0.42, *p*=.741). Furthermore, there were no general group effects (main effect group: *H*_*ER-A*_(3)=1.23, *p*=.732 and *H*_*ER-T*_(3)=2.20, *p*=.533, respectively), but participants became generally faster in correctly identifying facial expressions (main effect time: *F*(3,84)=46.13, *p*<.001, *η*^*2*^=.35), whilst accuracy did not change (*F*(3,84)=1.23, *p*=.270).

### Patterns of change in progestin profiles and RSFC are associated with differences in positive mood, but not any other mood-related or emotion recognition measures

Synchrony between pattern of changes in estrogenic and progestogenic hormonal profiles and parcelwise RSFC with changes in behavioral measures (i.e., ΔMood+ and ΔMood-, ΔDepression, ΔER-A and ΔER-T), respectively, were analyzed with IS-RSAs. For positive mood, the IS-RSA with change patterns in estrogen showed no significant association (*rho*(85)=0.12, *p*=.058), while the interindividual variation of changes in progestogens was resembled by positive mood patterns (*rho*(83)=0.20, *p*=.008). Fittingly, the change patterns of positive mood were also positively reflected by one of the hormone-modulated parcels, namely the right OFG (*rho*(85)=0.23, *p*=.036, see Figure 3D). In addition to Figure 3D, Table 2 lists all other frontal, subcortical, occipital, and cerebellar regions for which patterns of change in RSFC were significantly correlated to changes in positive mood scores. Patterns of changes in negative mood and depressive symptoms were not resembled by hormonal change (estrogens: *rho*_*Mood-*_(85)=0.04, *p*=.309 and *rho*_*Depression*_(86)=-0.004, *p*=.505, progestogens: *rho*_*Mood-*_(83)=-0.03, *p*=.617 and *rho*_*Depression*_(84)=-0.01, *p*=.531) nor by parcelwise RSFC patterns (ΔMood-: all *rho*(85)≤0.11, all *p*≥.689 and ΔDepression: all *rho*(88)≤0.15, all *p*≥.0.464).

Furthermore, interindividual variability of changes in emotion recognition accuracy or response times was also not reflected by the patterns of change in hormonal levels (estrogens: all *rho*(86)≤0.03, all *p*≥.352; progestogens: |*rho*(84)|≤0.06, all *p*≥.721) or parcelwise RSFC (ΔER-A: all *rho(88)*≤0.20, all *p*≥.197 and ΔER-T: all *rho(88)*≤0.08, all *p*≥.797).

## Discussion

We investigated possible neuroendocrinological modulations of brain architecture underlying changes in mood and emotion cognition in a heterogeneous sample of women with both relatively stable and drastically altered hormonal milieus as a result of OC intake. In line with the distribution of sex hormone receptors in the brain[2, 3], hormonal profile changes were reflected in the RSFC of a variety of predominantly frontal, subcortical, and cerebellar regions. However, these correlations between interindividual change patterns were only observed for endogenous and synthetic progestogens, but not for estrogens. Moreover, changes in positive mood resembled progestogen fluctuations as well as changes of RSFC of the progestogen modulated right OFG, suggesting a possible progestogen-induced neuroendocrinological modulation of positive mood. Changes in negative mood, depressive symptoms, as well as emotion recognition performance, showed no association with hormone concentration changes. However, OC discontinuation led to a reduction in depressive symptoms, while OC start showed the opposite pattern. Ultimately, these findings may provide better insight into possible underlying progestogen-related neuroendocrinological modulations of socio-affective processes.

The progestogen-related changes in RSFC in regions of high hormone receptor density, including the left hippocampus and brainstem, right (superior) orbitofrontal gyrus (OFG) and left middle/inferior frontal gyrus (MFG/IFG), and several regions in the right cerebellum, may be due to hormonally modulated genomic (i.e., involving gene transcription and protein synthesis) and non-genomic (i.e., extra-nuclear) signaling cascades[2, 3]. These signaling cascades can impact RSFC through processes such as synaptic pruning, neurite outgrowth, dendritic branching, myelination, and interactions with the neurotransmitter systems[5-8]. Through the proximity of these highly hormone innervated brain regions to multiple neurotransmitter pathways (e.g., serotonergic, GABAergic, glutamatergic, and dopaminergic)[2, 44], the hormone-modulated signaling cascades and direct modulation of the neurotransmitter pathways might also affect the RSFC of more distant brain regions. In this study, we for instance found changes in progestogen concentration resembling alterations in the RSFC of the left postcentral gyrus (PoCG) and parts of FFG/ITG, which are typically not characterized as high-density hormone receptor areas. Among the reported progestogen-associated brain regions, particularly the MFG and hippocampus have been repeatedly implicated to be affected by sex hormone concentrations[21, 45] and associated with potential alterations in socio-affective processing. For instance, MFG RSFC has been linked to the severity of mood problems in women suffering from premenstrual syndrome (PMS)[46]. Furthermore, in healthy women, MFG/IFG activity also appears to be modulated by sex hormones in terms of emotion regulation of negative affect[47] and face processing[48]. The hippocampus, on the other hand, is thought to play an important role in incorporating cognition (e.g., mentalizing) into emotional processing, for example in the case of social emotions[49, 50]. In addition, the cerebellum, which also has a high density of sex hormone receptors, has recently received more attention for its multifaceted role in socio-affective processing[51]. Notably, most of the progestogen-related regions have been implicated in socio-affective processing including mood and emotion recognition[3, 51, 52]. Therefore, the altered RSFC may be indicative of changes in socio-affective processing.

In contrast to changes in progestogens, changes in endogenous and synthetic estrogens were not reflected in RSFC change patterns of any investigated brain region. This is somewhat surprising as other studies found estrogens to be modulators of the dopaminergic[53] and the serotonergic neurotransmitter systems[54], which in turn implies that estrogens likely affect functional connectivity[55]. However, the lack of estrogen modulations on RSFC in our study might be explained by possible specificities of the estrogen-RSFC relationship: Previous studies found RSFC to be associated with estradiol in phases of rapidly changing hormonal levels (i.e., periovulatory phase) and network-based RSFC[42, 56], whereas our study investigates effects of endogenous and synthetic estrogen on a whole-brain level. Moreover, this study included only NC women in the early follicular phase, thus E2 levels were generally low and stable.

At the behavioral level, progesterone has been more consistently linked to emotion processing than estradiol[57]. In the present study, changes in the progestogen profiles were significantly correlated with positive mood alterations, while there were no mean-based differences in positive mood among women with different hormone profiles. A possible neuroendocrine mechanism of progestogen-induced mood alteration could pose the RSFC of the right OFG with its relevance for mood regulation and reward processing[58-60]. Moreover, in a directed analysis of hormone-modulated parcels (see Table S2 in supplement), change patterns in positive mood were also reflected by the RSFC in many of the progestogen-related brain regions including the left MFG/IFG, hippocampus, and right cerebellar regions, which were also partly in line with regions identified in a meta-analysis as neural substrates of positive and negative affect processing[61]. For negative mood, however, no significant associations were detected, probably due to the low variation in negative affect scores in our healthy participants. Overall, progestogens appear to modulate positive mood, which is in line with a previous longitudinal study reporting reduced positive mood after OC start compared to a naturally cycling control group[28].

Depression is a related construct of negative mood. In addition to negative mood, depressive symptoms encompass non-mood-related somatic and vegetative symptoms. In our study, the increase in self-reported depressive symptoms after the start of OC intake was mainly driven by single individuals, which is well in line with previous literature suggesting that merely a subsample of hormone-sensitive OC users could be susceptible to adverse mood effects[32]. Unlike OC start, the pattern in depressive scores was largely one-directional across OC discontinuers. Women who stopped OC intake had significantly higher depressive symptoms before OC discontinuation than women who continued OC intake. Therefore, a possible explanation for the found decrease in depressive symptoms in OC discontinuers could be that especially hormone-sensitive women, who experience OC-related side effects on mood, are more likely to discontinue OC intake than women without these side effects[62, 63]. However, only eight out of 25 OC discontinuers brought up mood problems as a reason for OC discontinuation. Taken together, in terms of self-reported depressive symptoms women seem to benefit from OC discontinuation although mood-related side effects were rarely mentioned as the reason for discontinuation. Although depressive symptoms changed significantly after initiation and discontinuation of OC intake, interindividual variation of changes in depressive symptoms was not reflected by patterns of changes in hormone profiles nor parcelwise RSFC. A possible explanation for this lack of (linear) association is the highly variable, non-linear effect of progestogens on the GABAergic system, which plays an important role in the neurobiological basis of depressive mood[64].

Due to its importance in daily life, several studies have investigated OC-related differences in emotion recognition[33]. Whilst most studies report altered emotion recognition in OC users[65-68], others did not find any significant difference in comparison to NC women[69, 70]. However, all these studies were cross-sectional and thus do not permit the inference of causal relationships. In our longitudinal study, we found no evidence of an OC-related or hormone profile-related effect on overall emotion recognition independent of the type of emotion displayed. Consequently, overall facial emotion recognition seems to be largely unaffected by changes in OC status and related changes in hormonal profiles. Nevertheless, it remains to be clarified whether the effects of OC on emotion recognition are emotion-dependent (as suggested by multiple studies, see for review: Gamsakhurdashvili, Antov [33]) or mediated by other factors such as progestin type, a more complex combination of different hormones, and/or individual susceptibilities through specific genotypes[66, 71].

While the current study presents new insights using a longitudinal design and the inclusion of OC discontinuers, several limitations ought to be addressed in future research. Firstly, the small sample size of OC starters limits statistical power to detect effects of OC initiation on mood and emotion recognition measures. Secondly, OC formulations were heterogenous with regards to dosages and type of progestins (i.e., androgenic vs antiandrogenic) used. Although we accounted for differences in face dosage by determining the bioavailability of synthetic hormones in the blood, we did not consider the different pharmacokinetic, pharmacodynamic and pharmacological properties of the progestins determining their biological potency. Preliminary findings suggest that androgenicity of progestins affects brain structure[72], likelihood of mood-related side effects[73] and emotion recognition performance[74]. Furthermore, the relation between RSFC and emotion recognition seems to be differently modulated by OC androgenicity[36]. Therefore, controlling for progestin types, larger sample sizes for stratification or coding of the estrogenic, progestogenic and androgenic properties to reflect biological potencies using information from in vivo bioassays and/or receptor studies[75] is recommendable to account more systematically for OC-related side-effects on the brain as well as socio-affective functions. Lastly, we focused on parcelwise RSFC as a first explorative step. Therefore, our analyses regarding neuroendocrinological modulations are limited to parcelwise RSFC. Functional networks, however, are often implicated as neural substrates of socio-affective processing[76]. Therefore, network-related functional connectivity may be a valuable complementation for identifying possible sex hormone-modulated neural substrates of mood and emotion recognition. Overall, large-scale dense-sampling longitudinal studies are needed in the future to enable well-defined analyses considering characteristics such as pill and progestin types, lifetime OC intake, reported OC side effects, genotypes, or other factors potentially moderating OC-related effects on neural functioning and socio-affective processes.

In summary, in a longitudinal design that included women with stable OC status (i.e., either with or without OC intake) and women with alternating OC status (i.e., initiation or cessation of OC intake), patterns of change in endogenous and synthetic progestogens resembled changes in RSFC of several frontoparietal, subcortical, and cerebellar regions implicated in socio-affective processing, including the left hippocampus, left MFG/IFG, right OFG, and right cerebellum. In particular, the right OFG, a region known for its implication in mood regulation and reward processing, was identified as a possible neural substrate involved in progestogen-related modulations of positive mood. Negative mood and depressive symptoms, on the other hand, were not directly associated with patterns in hormonal profiles or parcelwise RSFC; possibly due to the rather complex underlying progestogen-related mechanisms affecting the GABAergic system. Nevertheless, in direct hormone-independent comparisons of OC starters and discontinuers, active OC intake was associated with increased depressive symptoms. In OC starters this effect, however, may have been driven by individual hormone-susceptible women. Our results suggest that mood rather than emotion recognition is associated with hormonal profile changes and OC status. Nevertheless, future large-scale longitudinal studies with well-characterized samples are needed to systematically disentangle potential OC-related effects on brain function and structure, as well as their effects on socio-affective processes that are crucial for mental health and social functioning. A better understanding of OC-related effects on socio-affective processes and their neuroendocrinological substrates is necessary to respond to the growing unease of women not knowing about OC-related mental side effects, and ultimately to provide them with the knowledge they need to make well-informed contraceptive choices.

## Materials and Methods

### Participants

In total 104 healthy women between the age of 18 and 33 years were recruited through advertisements at the University of Tübingen, the University Hospital Tübingen as well as gynecological practices located in Tübingen and social media. However, 16 women dropped out before completing the study or were excluded from analysis due to time constraints (n=9), medical issues (n=6), or intrauterine device insertion (n=1). The women in the final sample (n=88, *m*_age_=23.7±3.4yrs) can be described by the following four hormonal profiles with the first two being relatively stable across measurements, whereas the last two are characterized by hormonal changes:

- Naturally cycling (NC) women (>6 months without the use of hormonal contraception) measured twice in a period of 4.8±1.2mths during the early follicular phase (day 2-5 of menstrual bleeding, **fNC women**: *n*=26, *m*_age_=23.8±3.5yrs).
- Long-term combined OC users (>6 months) measured twice in a period of 4.3±0.7mths during the active intake phase (day 3-21 of blister; **OC users**: *n*=26, *m*_age_=23.8±2.9yrs).
- OC discontinuers after long-term combined OC intake (>4 months) measured first during the one of the last two cycles of OC intake and after 5.0±1.1mths during the early follicular phase (day 2-5 of menstrual bleeding, >3 months NC, **OC discontinuers (dOC)**: *n*=25, *m*_age_=24.0±3.8yrs). Two OC discontinuers were measured outside the early follicular phase due to cycle irregularity/amenorrhea. One of them had her menstruation 2 days after the measurement.
- Combined OC starters (>4 months no hormonal contraception) measured first during the early follicular phase (day 1-5 of menstrual bleeding) before starting OC intake and 4.3±0.6mths later at the active OC intake (day 3-21 of blister, >3 months OC intake, **OC starters (sOC)**: *n*=11, *m*_age_=22.1±3.3yrs).

In general, to be eligible for study inclusion women had to be between 18 and 35 years old and have no history of any gynecological, mental, or neurological illnesses, pregnancies, nor any (other) hormonal or neuroactive medication (apart from L-thyroxine) in the past four months. The participants were comparable with respect to basic demographic and neuropsychological parameters including age, educational level, verbal intelligence, cognitive flexibility as well as depressive symptoms and trait anxiety. Furthermore, they reported similar thoughts and feelings experienced during the fMRI scan (see *Material and Methods* for information on the questionnaires and Table S1 for an overview of the sample descriptives). All women who entered the study as NC, were free of hormonal contraception for at least four months. Previous literature suggests that two to four months after contraceptive discontinuation the menstrual cycle should generally have normalized[77-79]. Only women taking or planning to take (i.e., OC starters) monophasic combined OCs were included in the study, with two exceptions in the OC discontinuers, who used a combined triphasic OC or the hormonal ring instead. A more detailed description of current and lifetime OC intake is provided in Table S1. The sample size is comparable to previous OC-related longitudinal studies[27, 28]. Furthermore, a-priori power analyses revealed that a group size of 12 should suffice to detect a medium group-by-time point effect at a power of .80 and α of .05 (G*Power 3.1.9.7). For IS-RSAs, there is no standard method of calculating power analyses. However, the similarity is calculated across 3486-3828 pairwise comparisons (i.e., 84×83/2 to 88×87/2) instead of 84 to 88 subjectwise observations as in other standard tests, thereby increasing statistical power significantly. Lastly, all women gave their written informed consent before participation. The study was approved by the Ethics committee of the Medical Faculty of the University of Tübingen (331/2016BO2).

### Procedure

At an initial screening appointment, potential exclusion criteria were monitored in a semi-structured interview, which accessed MRI compatibility, history of mental illness, and gynecological parameters (e.g., pregnancies, cycle irregularities, medication, and gynecological disorders). General mental health was assessed using the Structural Clinical Interview Screening (SCID screening)[80]; premenstrual symptoms were monitored using the Premenstrual Symptoms Screening Tool (PSST)[81]. Furthermore, we collected general information on sociodemographic characteristics (e.g., age and educational level) and neuropsychological functioning. Cognitive flexibility was assessed with the Trail Making Test (TMT)[82], whilst the Wortschatztest (WST)[83] measured crystallized verbal intelligence.Eligible participants were scheduled for two measurement appointments in an interval of three to eight months during the respective hormonal phase the women were assigned to (i.e., active OC intake day 3-21 or day 1-5 of the menstrual cycle, i.e., early follicular phase). Both measurement sessions followed the same procedure: Firstly, the participants performed the short Vienna Emotion Recognition Task (VERT)[84], before blood samples (i.e., 2×9ml EDTA) were drawn by medically trained staff for the plasma hormone determination. Subsequently, the participants were presented with a battery of questionnaires addressing mood with the Positive and Negative Affect Scale (PANAS)[85] and the Beck’s Depression Inventory II (BDI-II)[86]. Upon completion, participants were briefed on the MR scanner. To reduce head movements, participants were fixated with foam pads at the sides of their heads and adhesive medical tape across the forehead[87]. The (f)MRI measurement started with an anatomical scan (~8min) and was followed by a resting-state (RS) scan (~8min). The Amsterdam Resting-State Questionnaire[88] was carried out via the scanner’s intercom system to capture thoughts and feelings of the participants during the RS measurement. This was followed by three functional task-related measurements, which will be reported elsewhere[e.g., cross-sectional empathy: 89]. Total duration of each measurement session was about 3.5hrs. Figure 1 provides an illustration of the study design.

### Materials and Measures

#### Hormone sampling and assessment

After blood sampling, all samples were centrifuged for 15min at 4400rpm for the plasma extraction. The plasma samples were stored at -70°C. Estradiol (E2), ethinylestradiol (EE), progesterone (P4), progestin (P; i.e., dienogest, levonorgestrel, nomegestrol or chlormadinone acetate), and testosterone (T) concentrations were assayed with liquid chromatography-tandem mass spectrometry (LC-MS/MS) by the Institute of Pharmaceutical Science of the University Tübingen. A 1290 Infinity II UHPLC (Agilent Technologies, Germany) coupled with a QTRAP 4500 mass spectrometer (Sciex, USA) was used for analysis by selected reaction monitoring acquisition. Quality control standards and calibrants were included in the analysis batches of the samples. The analysis as well as the precision and accuracy levels (see supplementary material section S2.1, Table S3) fulfilled FDA bioanalytical method validation standards.

#### Acquisition of anatomical and functional MR images

Anatomical and functional MR scans were acquired at a 3T MR scanner (Siemens; Erlangen, Germany) using a 64-channel head coil. The anatomical image (i.e., T1-weighted MP2RAGE: 176 sagittal slices, TR/TE=5000/2.98ms, TI=700/2500ms, resolution=1×1×1mm, field of view (FOV)=256mm, flip angle [FA]=4/5°) helped in aligning the measurement of the functional scans to the anterior-posterior (AC-PC) axis and was used for coregistration during pre-processing. Whole-brain functional imaging was carried out with a multiband, T2-weighted EPI (i.e., echo planar imaging) sequence (slices=69, multiband factor=3, TR/TE=1500/34ms, resolution=2×2×2mm, FOV=192, FA=70°). To adjust for magnetic field deformations, a T2*-weighted field map (sequence: slices = 36, TR=400ms, TE=4.92/7.38ms, resolution=3.0×3.0×3.2mm, FOV=192mm, FA=60°) was run prior to EPI scans.

#### Emotion Recognition Task (VERT-K)

The short version of the Vienna Emotion Recognition Task (VERT-K) has been repeatedly used to successfully investigate emotion recognition performance under different ovarian hormone concentrations[69, 84, 90-92]. Participants are presented with a face stimulus showing one of the five basic emotions (happiness, sadness, anger, fear, and disgust) or a neutral expression. Next to the target face stimulus, participants can select the correct response among six response options to continue to the next trial. The written response options are presented in a randomized order for each trial. In total, stimuli consisted of 36 coloured pictures of European-American faces (i.e., 6 items per condition) balanced for actor sex. Intertrial intervals were 1s and the order of stimulus presentation was pseudo-randomized for emotional condition as well as the sex of the actor. The VERT-K lasts about two to four minutes.

#### Mood Measures

Positive and negative affective states were assessed by the previously validated Positive and Negative Affect Scale (PANAS)[85]. To assess their present mood, participants indicate on a 5-point Likert scale from ‘very little or not at all’ to ‘very much’, to what degree they experience an affective state described by the respective item at this very moment. The positive affect scale includes items such as happiness, enthusiasm, and alertness, whereas on the negative affect scale participants rate items like sadness, nervousness, and guilt. Each scale contains 10 different affective state items.

The Beck’s Depression Inventory II (BDI-II)[86] measures the degree of depressive symptomatology according to DSM-5 criteria during the past days to weeks. The BDI-II comprises 21 questions with four to six answer options. It covers symptoms including restlessness, self-esteem, loss of energy, difficulty concentrating, sleep and appetite disturbances as well as changes in body image and weight.

### Statistical Analysis

Statistical analyses of sample characteristics, hormone and behavioral measures were conducted in *IBM SPSS Statistics 25*. Representational similarity analyses (RSA) were carried out with custom *Python V3*.*8*.*8* scripts using Jupyter Notebook (anaconda3). The applied statistical threshold is α=.05 and all post-hoc tests and planned contrasts were Bonferroni-corrected. RSA was corrected for multiple comparisons using permutation testing.

#### Sample characteristics

Information on sample characteristics such as age, education, neuropsychological as well as OC-related aspects and their analyses are reported in the supplementary material (section S1 as well as Table S1 and section S2.2, respectively).

#### Group differences and changes across time points in sex hormones, emotion recognition, and mood measures

Group differences at the two measurement time points and in the change over time were analysed using 4 (between: group – fNC, OC, dOC, and sOC) by 2 (within: time – T0 and T1) mixed ANOVAs. This model was run on endogenous and exogenous sex hormones (i.e., E2, P4, EE, and P, respectively), emotion recognition (i.e., overall accuracy in % and average response times of correct trials in s), and mood measures including the mean scores of the positive and negative PANAS scales, respectively, as well as the total score of depressive symptoms measured by the BDI-II. Due to the relatively large robustness of ANOVA to non-normally distributed data[93], we decided to use mixed ANOVAs also for dependent variables with non-normally distributed residuals. However, post-hoc tests were chosen dependent on the data’s normality and homogeneity including non-parametric tests such as Kruskal Wallis and Wilcoxon tests. If the data was normally distributed but homogeneity was violated Welch’s ANOVA and Games-Howell testing was chosen. Post-hoc analyses were corrected for multiple comparisons using Bonferroni correction.

#### Preprocessing of RS fMRI data and computation of parcelwise RSFC

##### Preprocessing of anatomical and functional (f)MRI scans

For preprocessing of all scans, the automated preprocessing pipeline fMRIprep 20.2.3 was used[94, 95]. For a detailed description of all fMRIprep preprocessing steps please refer to the supplementary material section S2.2. Anatomical scans were corrected for intensity non-uniformity and subsequently skull stripped and segmented in cerebral spinal fluid (CSF), white matter (WM), and grey matter (GM). Finally, the anatomical image was normalized to the Montreal Neurological Institute space (MNI152NLin6Asym[96]). Functional scans underwent a susceptibility distortion correction using field maps, coregistration to the anatomical scan, realignment, slice time correction, and normalisation to the MNI152NLin6Asym template. Framewise displacement was lower than 2mm for all scans. The output was passed on to custom Matlab scripts for the computation of parcelwise RSFC.

##### Computation of parcelwise RSFC

A parcelwise approach was chosen to extract activity time courses to lower the dimensionality of data and reduce computational costs and slight misalignments across participants. Considering that particularly subcortical regions and the cerebellum are densely packed with hormone receptors[2], the functional parcellation of the Shen atlas[39] seemed a suitable choice for this study, as unlike other parcellations it includes subcortical and cerebellar parcels. Furthermore, with its 268 functional parcels, the Shen parcellation provides good spatial specificity. For each time point, the parcelwise extracted activation time course was adjusted by the exclusion of the variance belonging to possible confounders such as mean WM, CSF, and global signal. Subsequently, the parcelwise time series was computed as the averaged first eigenvariate of all constituting voxels within a given parcel. Parcelwise connectivity was computed as Fisher’s Z scores of the pairwise Pearson’s correlations between each parcel’s time series with those of all other parcels. This computation was done for each subject and separately for each of the two measurement time points. The resulting subjectwise connectivity matrices were subsequently used to compute representative distance matrices between the subjects, which in turn were utilized for intersubject representational similarity analyses (RSAs).

#### Intersubject representational similarity analyses (IS-RSA) with changes in parcelwise whole-brain RSFC, hormone profiles, and socio-affective measures

To investigate the potential relationships between interindividual variability in patterns of change functional connectivity profiles, hormones, and socio-affective measures, we computed IS-RSAs. An overview of the different IS-RSAs computed in this study is provided in Figure 2B. IS-RSA is a computational technique based on pairwise comparisons of subjects’ responses to reveal their representation in a higher order space. Computing IS-RSA includes two main steps: (1) calculation of pairwise (subject-by-subject) similarity matrices (i.e., representational distance matrices) for each measurement modality and (2) comparison of two similarity matrices using a Mantel test.

##### Representational distance matrices (RDMs)

An IS-RDM is a two-dimensional array containing the subject-by-subject pairwise distances between the elements of a measure’s set. Euclidean distance between subjects was used to compute the distances between subjects with regards to changes (T1 minus T0) of a parcel’s averaged RSFC with all other parcels (n=268), whereas standardised Euclidean distance was used for the remaining RDMs to assure comparability among dimensions within an RDM but also between different RDMs. RDMs were always calculated from the change scores (Δ: T1 minus T0) in multiple-dimensional spaces (i.e.: estrogens: Δ of E2 and EE concentrations, progestogens: Δ of P4 and P concentrations, RSFC: Δ of RSFC with every other parcel, positive mood: Δ of each item in Positive Affect Scale, negative mood: Δ of each item in Negative Affect Scale, Emotion Recognition Accuracy: Δ correct yes/no per item, Emotion Recognition Time: Δ Average response time of correct items per emotion). Emotion recognition response times were averaged across emotions otherwise many trials would be excluded as response times were only used for correctly solved trials. An itemwise over mean-based approach was chosen to preserve individual variation[97].

##### Representational similarity analyses (RSAs)

RDMs of different modalities were compared using Spearman rank correlation as a Mantel test (see Figure 2B). All *p*-values were computed using permutation-based testing. To this end, hormonal or behavioral RDMs were permutated in 10,000 iterations and a null distribution of 10,000 surrogate correlation values was built. The corrected p-value was obtained by locating the observed Spearman Rho coefficient of the true relationships on the created null distribution. For IS-RSAs involving the parcelwise RDMs, *p*-values were additionally controlled for multiple comparisons by creating a family-wise null distribution of 10’000 correlation coefficients using the maximum value per iteration of the 268 null distributions. Since we were especially interested in whether hormone-associated RSFC change is associated with changes in socio-affective measures, we ran additional IS-RSAs with only parcels in which RSFC was associated with hormone profile changes (n=12). For these analyses, p-values were corrected for 12 parcels only (see Table S2 in supplementary material).

## Authors contributions

**Ann-Christin S. Kimmig:** Conceptualization, Methodology, Project administration, Investigation, Data Curation, Formal analysis, Visualisation, Writing – original draft, Writing – review & editing. **Patrick Friedrich:** Conceptualization, Methodology, Data Curation, Formal analysis, Visualisation, Writing - review & editing. **Bernhard Drotleff:** Validation, Formal analysis (Hormones), Data curation, Writing - review & editing. **Michael Lämmerhofer:** Resources, Supervision (Hormones), Writing - review & editing. **Inger Sundström-Poromaa:** Methodology, Writing - review & editing. **Susanne Weis:** Conceptualization, Methodology, Supervision (Analyses), Resources, Writing – review & editing. **Birgit Derntl:** Conceptualization, Methodology, Project administration, Supervision, Funding acquisition, Resources, Writing – review & editing

## Acknowledgements

We would like to thank Annika Birrenbach, Anna Gärtner, Melina Grahlow, Leonie Matkei and Theresa Schell for their assistance during participant recruitment and data collection.

