## Supplementary Material for "To start or to discontinue the pill – changes in progestogens reflected by resting-state connectivity and positive mood"

### S1. Results

**Sample characteristics.** Table S1 displays means and standard deviations for sociodemographic (i.e., age and educational level), handedness, neurocognitive (i.e., verbal intelligence and cognitive flexibility), OC-related (i.e., lifetime and current OC intake duration, age of onset, type of current OC and duration since last OC intake) and resting-state related (8 subscales of the Amsterdam Resting State Questionnaire) data. The different groups of women did not differ in any sociodemographic and neurocognitive characteristics (all |*H*|= 2.97, *p*≥*.396*). However, the interval between the measurement points was significantly larger in OC discontinuers compared to OC users with continued intake (*p=.034*). The longer interval for OC discontinuers is largely attributable to the COVID19 pandemic, as measurements had to be paused for several months. The measurements of the OC group were finished before the outbreak of COVID19.

The onset age of OC intake in women who had prior experience with OC did not differ across groups (*F*(3,65)=1.35, *p*=.265). However, the lifetime exposure to OCs was significantly larger in current OC users and OC discontinuers than in naturally cycling women (*H*(3)=17.61, *p*=.001; dOC, OC>fNC: all *p* ≤ .003). OC users and OC discontinuers did not differ in the current OC intake duration (*t*(1,49)=0.09, *p*=.762) and OC androgenicity was similarly distributed among the groups with active OC intake during the study (*p*=.699).

None of the eight ARSQ scales showed an interaction between group and time (all |*F*|≤1.78, all *p*≥.158). A main effect of group was found for somatic awareness (*F*(3,83)=3.06, *p*=.033, $\eta^2$=.10), however, Bonferroni corrected post-hoc test did not reveal significant pairwise differences (all *p*≥*.066*). Across all groups, there was a significant decrease in somatic awareness (*F*(3,83)=7.34, *p*=.008, $\eta^2$=.08) and theory of mind (*F*(3,83)=4.17, *p*=.044, $\eta^2$=.05) from the first to the second measurement (*F*(3,83)=3.06, *p*=.033, $\eta^2$=.10). However, when correcting for multiple comparison across all eight scales, none of these findings remain significant.

*Table S1. Sample description (mean and standard deviation if not otherwise specified) including history of OC intake and information on thoughts and feelings during the resting-state scan measured by the Amsterdam Resting State Questionnaire*

| | **fNC** | **OC** | **dOC** | **sOC** | **$p_{group}$** |
|---|---|---|---|---|---|
| **N** | 26 | 26 | 25 | 11 | |
| Age (years) | 23.8 (3.5) | 23.8 (3.0) | 23.9 (3.9) | 22.2 (3.3) | .396 |
| Education (l/m/h) | 0/18/8 | 0/17/9 | 2/15/8 | 0/8/3 | .775 |
| Edinburgh-Handedness Score | 91.6 (39.2) | 91.6 (27.2) | 96.8 (9.5) | 63.6 (76.7) | .396 |
| WST verbal IQ | 107 | 105 | 105 | 102 | .584 |

| | | | | | | |
|---|---|---|---|---|---|---|
| WST raw scores | | 32.9 (3.5) | 32.4 (2.2) | 32.3 (2.6) | 31.3 (4.1) | |
| Cognitive flexibility (TMTB-A; sec) | | 17.0 (9.5) | 17.5 (10.4) | 19.1 (9.6) | 17.3 (11.1) | .678 |
| Time between T0 and T1 (months) | | 4.8 (1.2) | 4.3 (0.7) | 5.1 (1.1) | 4.3 (0.6) | **.026** *dOC > OC* |
| **OC-related information** | | | | | | |
| Lifetime OC intake (yes/no) | | 15/11 | 26/0 | 25/0 | 7/4 | |
| Age of OC intake onset | | 17.4 (1.7) | 16.8 (1.6) | 17.7 (2.4) | 16.3 (1.6) | .265 |
| Lifetime OC intake duration (months) | | 31.3 (45.0) | 72.1 (42.9) | 68.4 (34.3) | 41.9 (50.4) | **.001** *dOC, OC > fNC* |
| Current OC intake (months) | | | 57.8 (34.1) | 60.7 (32.7) | | .762 |
| Pill Androgenicity (A/AA) | | | 15/11 | 14/11 | 8/3 | .699 |
| 0.03mg EE/2mg Dienogest | | | 10 | 9 | 3 | |
| 0.015 EE/0.12 Etonogestrel (Nuvaring) | | | | 1 | | |
| 0.03mg EE/2mg Chlormadinone acetate | | | 1 | 2 | | |
| 0.03mg EE/0.15mg Levonorgestrel | | | 3 | | 5 | |
| 0.03mg EE/0.125mg Levonorgestrel | | | 3 | 2 | 1 | |
| 0.03-0.04mg EE/0.05-0.125 mg Levonorgestrel | | | | 1 | | |
| 0.2mg EE/0.1mg Levonorgestrel | | | 9 | 8 | 1 | |
| 0.2mg EE/0.15mg Desogestrel | | | | 1 | 1 | |
| 0.35mg EE/0.25mg Norgestimate | | | | 1 | | |
| Duration since last OC intake (months) | | 27.4 (26.0) | | | 17.5 (19.4) | .158 |
| **Amsterdam Resting State Questionnaire (ARSQ)*** | | | | | | |
| Discontinuity | T0 | 2.7 (0.5) | 2.3 (0.6) | 2.5 (0.7) | 2.6 (0.8) | 1.000 |
| | T1 | 2.7 (0.5) | 2.6 (0.7) | 2.4 (0.7) | 2.7 (0.9) | 1.000 |
| | $p_{T0T1}$ | 1.000 | .084 | 1.000 | 1.000 | |
| Theory of Mind | T0 | 3.5 (0.9) | 3.3 (0.8) | 3.3 (0.6) | 3.5 (0.6) | 1.000 |
| | T1 | 3.4 (1.0) | 3.1 (0.8) | 3.5 (0.6) | 3.0 (0.9) | .798 |
| | $p_{T0T1}$ | 1.000 | .552 | 1.000 | .666 | |
| Self | T0 | 3.4 (0.9) | 2.8 (0.8) | 3.0 (0.8) | 3.1 (0.9) | .660 |
| | T1 | 3.3 (0.9) | 2.9 (0.9) | 3.3 (0.8) | 3.2 (0.9) | 1.000 |
| | $p_{T0T1}$ | 1.000 | 1.000 | .774 | 1.000 | |

| | | | | | | |
|---|---|---|---|---|---|---|
| Planning | T0 | 3.2 (0.9) | 2.7 (0.8) | 3.00 (0.5) | 3.0 (0.7) | .990 |
| | T1 | 2.9 (0.9) | 2.9 (0.8) | 3.2 (0.6) | 3.0 (0.8) | 1.000 |
| | $p_{T0T1}$ | .768 | 1.000 | 1.000 | 1.000 | |
| Sleepiness | T0 | 2.7 (1.2) | 2.5 (1.1) | 2.9 (1.0) | 2.6 (1.2) | 1.000 |
| | T1 | 3.1 (0.9) | 3.0 (1.1) | 2.8 (0.9) | 2.9 (1.2) | 1.000 |
| | $p_{T0T1}$ | .456 | .168 | 1.000 | 1.000 | |
| Comfort | T0 | 3.6 (0.8) | 3.6 (0.7) | 3.4 (1.0) | 3.0 (1.1) | 1.000 |
| | T1 | 3.7 (0.8) | 3.6 (0.8) | 3.8 (0.8) | 3.1 (0.7) | .300 |
| | $p_{T0T1}$ | 1.000 | 1.000 | .588 | 1.000 | |
| Somatic Awareness | T0 | 2.8 (0.9) | 2.7 (0.9) | 3.4 (1.0) | 2.7 (0.9) | .372 |
| | T1 | 2.5 (0.9) | 2.5 (0.9) | 3.1 (1.1) | 2.3 (1.0) | .576 |
| | $p_{T0T1}$ | 1.000 | .648 | 1.000 | .348 | |
| Health concerns | T0 | 1.5 (0.5) | 1.3 (0.4) | 1.4 (0.5) | 1.3 (0.5) | 1.000 |
| | T1 | 1.4 (0.4) | 1.4 (0.5) | 1.4 (0.4) | 1.2 (0.3) | 1.000 |
| | $p_{T0T1}$ | 1.000 | 1.000 | 1.000 | 1.000 | |

Note: *For ARSQ, all depicted p-values have been corrected for multiple comparisons within the respective scale (i.e., $p_{Bonf}$=$p$*6). l=no higher education entrance qualification, m=higher education entrance qualification, h=university degree. T0 – first measurement, T1 – second measurement, fNC – early follicular naturally cycling women, OC – Oral Contraceptive users, dOC – OC users discontinuing OC intake after T0, sOC – early follicular naturally cycling women starting OC intake after T0, WST – Wortschatztest, IQ – intelligence quotient, TMTB-A – Difference between part B and A of the Trial Making Task, EE - Ethinylestradiol

**Endogenous E2.** The mixed ANOVA for E2 showed a significant group-by-time interaction ($F$(3,82)=25.00, $p$<.001, $\eta^2$=.48), and a main effect of group ($H$(3)=40.99, $p$<*.001*, post-hoc: fNC, dOC, sOC > OC, all $p$≤.019). The main effect of time ($F$(1,82)=0.525, $p$=.53) remained non-significant. Disentangling the interaction with Bonferroni correction revealed that at both time points (T0: $H$(3)=47.41, $p$<.001; T1: $H$(3)=51.79, $p$<.001, all pairwise comparisons: $p$≤.001) E2 was significantly higher in NC than in OC women (i.e., T0: fNC, sOC > OC, dOC; T1: fNC, dOC > OC, sOC).Furthermore, there was no significant change in E2 for women measured twice during the same hormonal status (all $p$≥.606). OC discontinuers had a significant increase of E2 across time ($Z$=-3.92, $p$<*.001*), whereas for OC starters E2 was significantly suppressed after start of OC intake ($Z$=-2.93, $p$=*.020*).

**Synthetic EE.** At T0, OC users and OC discontinuers did not differ significantly in their EE levels ($H$(1)=0.54, $p$=*.462*). As expected, at T1 no EE was traceable in the blood of OC discontinuers. However, OC starters had significantly higher blood serum levels of EE after about four months of OC intake compared to long-term OC users with an average intake duration of about five years ($H$(1)=13.48, $p$<*.001*). Furthermore, women with continued OC intake showed a significant decrease of EE concentration between the measurements ($Z$=-2.31, $p$=.021).

**Endogenous P4.** Analogous to endogenous E2, the mixed ANOVA on P4 concentrations revealed a significant group-by-time effect ($F$(3,83)=3.67, $p$=.015, $\eta^2$=.12), whilst there was no main effect of time ($F$(1,83)=1.05, $p$=.309). The main effect of group ($H$(3)=39.83, $p$<.001*;* post-hoc: fNC, dOC, sOC > OC, all $p$≤.004) again is explained by the interaction term. Naturally cycling women at T0 (i.e., fNC and sOC) had significantly higher P4 serum concentrations compared to OC users (i.e., OC and dOC; $H$(3)=46.40, $p$<.001, post hoc: all $p$<.001). For T1, only P4 of long-term OC users but not of OC starters was significantly lower than the concentrations in the naturally cycling women (*H*(3)=24.33, *p*<.001, post hoc: all *p*<.001). P4 increased significantly after discontinuation of OCs (*Z*=-3.71, *p=.001*). No other significant differences in P4 between the timepoints were found after Bonferroni correction (all *Z*≤2.58, all *p*≥.060).

**Synthetic P**. Synthetic P levels of OC discontinuers at T0 and OC starters at T1 were not significantly different from the OC group (all *H*≤0.89, all *p*≥.366). No time differences were found for synthetic P of women with continued OC intake (Z=-0.72, *p*=.469).

**Endogenous T.** The mixed ANOVA revealed a significant group-by-time interaction ($F(3,83)=4.61$, $p=.005$, $\eta^2=.14$). While there were no group differences at the individual measurement timepoints (all *H*≤4.34, all *p*≥1.000), T concentrations decreased significantly in the fNC group from T0 to T1 (*t*(25)=3.16, *p*=.024) with no further significant results in the other groups (all *p*≥.312). The main effects of group (*H*(3)=1.27, *p*=.737) and time (*F*(1,83)=1.23, *p*=.271) remained non-significant.

**Progestogen-related RSFC-behavior RSAs.** Since we were specifically interested in whether mood changes were reflected in changes in RSFC of parcels associated with progestogen change patterns, we ran analyses only including the parcels showing a hormone-association (i.e., permutation-based multiple comparison correction for 12 parcels). For positive mood, several parcels within the right cerebellum (i.e., Crus 2, VI, and IX), the right FFG/ITG, left hippocampus, and left MFG/IFG also show significant, positive associations in permutation-based IS-RSA (all *rho*(85)≥0.17, all *p*≤.047 [adjusted for 12 parcels], see Table S2). No other behavioural measure was resembled by parcelwise RSFC patterns of previously identified progestogen-related parcels (negative mood: all *rho*(85)≤0.09, all *p*≥.441, depressive symptoms: all *rho*(88)≤0.02, all *p*≥.774, emotion recognition accuracy: all *rho(88)*≤0.05, all *p*≥.630; emotion recognition response times: all |*rho(88)*|≤0.12, all *p*≥.900).

*Table S2.* Overview of significant IS-RSA results between change patterns of parcelwise RSFC of parcels revealed by progestogen IS-RSA and positive mood (corrected for multiple comparisons for 12 regions). Parcelwise MNI coordinates (centre of mass) and functional networks belonging are reported according to Finn et al., 2015[1].

| ***Shen Parcels*** | ***rho*** | ***MNI coordinates*** | | | ***Network*** | ***$p_{corr}$*** |
|---|---|---|---|---|---|---|
| | | *x* | *y* | *z* | | |
| ***Associated with progestogens (n=84 participants)*** | | | | | | |
| *R OFG* | 0.23 | 5.1 | 34.9 | -17.4 | Default Mode | .005 |
| *R Cerebellum Crus 2* | 0.20 | 11.7 | -84.1 | -34.6 | Frontoparietal | .022 |
| *R Cerebellum VI* | 0.19 | 21.1 | -54.8 | -23.8 | Subcortical-Cerebellar | .024 |
| *R FG/ITG* | 0.18 | 31.5 | 0.7 | -44.4 | Motor | .034 |
| *L Hippocampus* | 0.18 | -35.7 | -24.8 | -14.9 | Subcortical-Cerebellar | .036 |
| *L MFG/IFG* | 0.18 | -43.0 | 42.0 | 11.0 | Frontoparietal | .044 |
| *R Cerebellum IX* | 0.17 | 7.1 | -53.7 | -34.4 | Subcortical-Cerebellar | .047 |

*All p values are corrected for multiple comparisons by permutation-based testing. L – left, R – right, OFG – orbitofrontal gyrus, FFG/ITG – fusiform gyrus/inferior temporal gyrus, MFG/IFG – middle frontal gyrus/inferior frontal gyrus*

### S2. Methods

#### S2.1 Hormone assessment

*Table S3. Quality control of hormone assessment including the dynamic range, interday precision as well as accuracy for endogenous and synthetic sex hormones.*

| | Dynamic range (pg/mL) | Interday Precision (%) | Interday Accuracy (%) |
|---|---|---|---|
| **Endogenous Sex Hormones** | | | |
| Estradiol (E2) | 3.5-5179.1 | 7.0-9.1 | 96.8-100.5 |
| Progesterone (P4) | 1.0-47657.0 | 6.4-9.9 | 97.0-104.6 |
| Testosterone | 1.9-11438.0 | 7.4-9.9 | 94.3-106.5 |
| **Synthetic Sex Hormones** | | | |
| Ethinylestradiol (EE) | 2.0-3000.0 | 5.6-12.3 | 97.1-99.9 |
| Progestins* | 10.0-20000.0 | 4.4-11.1 | 93.4-109.2 |

Note: *In this table the minimum and maximum levels across all four progestins are reported as a collective

#### S2.2 Statistical Analyses

**Sample characteristics.** Cognitive flexibility was derived by subtracting the TMT-A from the TMT-B time[2]. Age, handedness, verbal intelligence, cognitive flexibility, the time between measurements, lifetime OC intake duration and duration since stopped OC intake were analysed using Kruskal Wallis tests, as their residuals were not normally distributed (determined by visual inspection and Shapiro Wilk tests: $p<.05$). Onset age of OC intake as well as current OC intake were analysed using an ANOVA with group as between factor (normality: yes, homogeneity of variances: yes). Education level and OC androgenicity was analyzed using the exact Fisher's test. For each ARSQ scale, a 4 (between: group) by 2 (within: timepoints) mixed ANOVA was performed. To determine p-values at the two different timepoints and differences between timepoints within groups, ANOVA (normality: yes, homogeneity of variances: yes), Welch ANOVA (normality: yes, homogeneity of variances: no), Kruskal Wallis tests (normality: no) and paired t-test (normality: yes) or Wilcoxon tests (normality: no) were used, respectively, dependent on the normality and homogeneity of the data. For all analyses, Bonferroni-corrected post-hoc tests and pairwise comparisons are reported for parametric as well as non-parametric tests.

**fMRI preprocessing**. The following sections on anatomical and functional preprocessing were automatically generated by fMRIprep[3] and were minimally adjusted with respect to formatting.

Results included in this manuscript come from preprocessing performed using fMRIPrep 20.2.3 (RRID:SCR_016216)[3, 4], which is based on Nipype 1.6.1 (RRID:SCR_002502)[5, 6].

**Anatomical data preprocessing**. A total of 2 T1-weighted (T1w) images were found within the input BIDS dataset. All of them were corrected for intensity non-uniformity (INU) with N4BiasFieldCorrection[7], distributed with ANTs 2.3.3 8 (RRID:SCR_004757)[8]. The T1w-reference was then skull-stripped with a Nipype implementation of the antsBrainExtraction.sh workflow (from ANTs), using OASIS30ANTs as target template. Brain tissue segmentation of cerebrospinal fluid (CSF), white-matter (WM) and gray-matter (GM) was performed on the brain-extracted T1w using fast (FSL 5.0.9, RRID:SCR_002823)[9]. A T1w-reference map was computed after registration of 2 T1w images (after INU-correction) using mri_robust_template (FreeSurfer 6.0.1)[10].

Brain surfaces were reconstructed using recon-all (FreeSurfer 6.0.1, RRID:SCR_001847)[11], and the brain mask estimated previously was refined with a custom variation of the method to reconcile ANTs-derived and FreeSurfer-derived segmentations of the cortical gray-matter of Mindboggle (RRID:SCR_002438)[12]. Volume-based spatial normalization to two standard spaces (MNI152NLin6Asym, MNI152NLin2009cAsym) was performed through nonlinear registration with antsRegistration (ANTs 2.3.3), using brain-extracted versions of both T1w reference and the T1w template. The following templates were selected for spatial normalization: FSL's MNI ICBM 152 non-linear 6th Generation Asymmetric Average Brain Stereotaxic Registration Model [Evans, Janke [13], RRID:SCR_002823; TemplateFlow ID: MNI152NLin6Asym], ICBM 152 Nonlinear Asymmetrical template version 2009c [Fonov, Evans [14], RRID:SCR_008796; TemplateFlow ID: MNI152NLin2009cAsym].

**Functional data preprocessing**. For each of the 2 BOLD runs found per subject (across all tasks and sessions), the following preprocessing was performed. First, a reference volume and its skull-stripped version were generated using a custom methodology of fMRIPrep. A B0-nonuniformity map (or fieldmap) was estimated based on a phase-difference map calculated with a dual-echo GRE (gradient-recall echo) sequence, processed with a custom workflow of SDCFlows inspired by the epidewarp.fsl script and further improvements in HCP Pipelines[15]. The fieldmap was then co-registered to the target EPI (echo-planar imaging) reference run and converted to a displacements field map (amenable to registration tools such as ANTs) with FSL's fugue and other SDCflows tools. Based on the estimated susceptibility distortion, a corrected EPI (echo-planar imaging) reference was calculated for a more accurate co-registration with the anatomical reference. The BOLD reference was then co-registered to the T1w reference using bbregister (FreeSurfer) which implements boundary-based registration[16]. Co-registration was configured with six degrees of freedom. Head-motion parameters with respect to the BOLD reference (transformation matrices, and six corresponding rotation and translation parameters) are estimated before any spatiotemporal filtering using mcflirt (FSL 5.0.9)[17]. BOLD runs were slice-time corrected using 3dTshift from AFNI 20160207 (RRID:SCR_005927)[18]. The BOLD time-series (including slice-timing correction when applied) were resampled onto their original, native space by applying a single, composite transform to correct for head-motion and susceptibility distortions. These resampled BOLD time-series will be referred to as preprocessed BOLD in original space, or just preprocessed BOLD. The BOLD time-series were resampled into standard space, generating a preprocessed BOLD run in MNI152NLin6Asym space. First, a reference volume and its skull-stripped version were generated using a custom methodology of fMRIPrep.

Several confounding time-series were calculated based on the preprocessed BOLD: framewise displacement (FD), DVARS and three region-wise global signals. FD was computed using two formulations following Power (absolute sum of relative motions, Power, Mitra [19]) and Jenkinson (relative root mean square displacement between affines, Jenkinson, Bannister [17]). FD and DVARS are calculated for each functional run, both using their implementations in Nipype (following the definitions by Power, Mitra [19]). The three global signals are extracted within the CSF, the WM, and the whole-brain masks. Additionally, a set of physiological regressors were extracted to allow for component-based noise correction (CompCor)[20]. Principal components are estimated after high-pass filtering the preprocessed BOLD time-series (using a discrete cosine filter with 128s cut-off) for the two CompCor variants: temporal (tCompCor) and anatomical (aCompCor). tCompCor components are then calculated from the top 2% variable voxels within the brain mask. For aCompCor, three probabilistic masks (CSF, WM and combined CSF+WM) are generated in anatomical space. The implementation differs from that of Behzadi et al. in that instead of eroding the masks by 2 pixels on BOLD space, the aCompCor masks are subtracted a mask of pixels that likely contain a volume fraction of GM. This mask is obtained by dilating a GM mask extracted from the FreeSurfer's aseg segmentation, and it ensures components are not extracted from voxels containing a minimal fraction of GM. Finally, these masks are resampled into BOLD space and binarized by thresholding at 0.99 (as in the original implementation). Components are also calculated separately within the WM and CSF masks. For each CompCor decomposition, the k components with the largest singular values are retained, such that the retained components' time series are sufficient to explain 50 percent of variance across the nuisance mask (CSF, WM, combined, or temporal). The remaining components are dropped from consideration.

The head-motion estimates calculated in the correction step were also placed within the corresponding confounds file. The confound time series derived from head motion estimates and global signals were expanded with the inclusion of temporal derivatives and quadratic terms for each[21]. Frames that exceeded a threshold of 0.5 mm FD or 1.5 standardised DVARS were annotated as motion outliers. All resamplings can be performed with a single interpolation step by composing all the pertinent transformations (i.e., head-motion transform matrices, susceptibility distortion correction when available, and co-registrations to anatomical and output spaces). Gridded (volumetric) resamplings were performed using antsApplyTransforms (ANTs), configured with Lanczos interpolation to minimize the smoothing effects of other kernels[22]. Non-gridded (surface) resamplings were performed using mri_vol2surf (FreeSurfer).

Many internal operations of fMRIPrep use Nilearn 0.6.2 (RRID:SCR_001362)[23], mostly within the functional processing workflow. For more details of the pipeline, see the section corresponding to workflow in *fMRIPrep's* documentation (https://fmriprep.readthedocs.io/en/latest/workflows.html).
